# A general decrease of normalized ncDNA in evolution

**DOI:** 10.1101/511196

**Authors:** Francisco Javier Lobo-Cabrera

## Abstract

Complexity is often associated with increasing non-coding DNA (ncDNA). For example, the human genome is in its vast majority ncDNA. Here, it is hypothesized that normalized ncDNA (nncDNA) has in fact diminished in evolution. This definition of ncDNA content takes into consideration total proteomic content. It is shown that by reducing their normalized ncDNA, organisms may have obtained more complexity in evolution. Also, a potential connection between ncDNA, proteome information and chromatin interactions in mice and humans is presented.

## Introduction

Life comprises a large number of forms; all of which composed of one or more cells [1]. In these, one or several DNA molecules carry the genetic information. DNA therefore controls cellular activities, and is transmitted across generations [1][2].

Initially, the quantity of DNA was thought to determine complexity in organisms [3]. Simpler organisms would have less cellular DNA than more complex ones. However, it soon became clear that it was not the case. Neither the total amount of DNA nor the number genes seem to correlate accurately with complexity. This has been termed the C-paradox [3] and the G-paradox [4], respectively.

A more successful approach to account for complexity consists of non-coding DNA (ncDNA). ncDNA is defined as DNA that does not encode proteins [5]. The percentage of ncDNA in the genome has been reported as a valid predictor of complexity [6][7]. In this way, it is not large amounts of DNA or number of genes, but rather large ncDNA percentages, what seems to characterize complex organisms [6][7].

Interestingly, ncDNA comprises i) identified regulatory sequences, but especially ii) large regions of unknown function. The latter are primarily transposon-derived DNA; i.e DNA created by molecular parasites known as transposable elements. As they lack of apparent function, these regions are often referred as *junk* DNA [8]. The fact that genomes of advanced species are mainly ncDNA is thus intriguing. So, even though ncDNA is an effective indicator of complexity, the reason for this remains unclear.

Another more intuitive predictor of complexity can be suggested. In this case, it is the total number of different proteins that an organism can display –referred here as Proteome Information Units (PIUS). The referred amount does not correspond to the number of genes, as alternative splicing and post-translational modifications allow genes to code for multiple proteins. Consequently, PIUS take into consideration the standard set of proteins plus their isoforms.

In the present work, the relation between these two complexity indicators --ncDNA percentage and Proteome Information Units-- is assessed. The ratio, named here normalized ncDNA (nncDNA), reveals a characteristic behavior which could provide insights into evolution.

## Results

### A general decrease in normalized ncDNA

Previous studies [6][7] were used to extract information about ncDNA content for multiple species. The focus was not on the total amount of kilobases of ncDNA, but on ncDNA percentages in relation to the whole genome.

On the other hand, Uniprot [9] served as a source of proteomic data. Particularly, reference proteomes were employed. Reference proteomes constitute a series of proteomes selected to represent biological diversity [9]. Uniprot provides a general overview of the data in terms of i) number of proteins of the haploid genome, ii) number of isoforms and iii) mappings. All of these elements are indicated for each species. Here, the elements i) and ii) were added in each case to calculate the PIUS values as discussed in the introduction.

Once the ncDNA data and proteomic data were collected, the next step was to select the organisms for which both types of information was available. The results include a total of 26 species, ranging from bacteria to mammals. Table 1 contains for each organism its correspondent i) ncDNA percentage, ii) PIUS, iii) ncDNA percentage ∙ PIUS⁻¹ ratio (i.e nncDNA) and iv) an identifying letter.

**Table 1.** ncDNA percentage, PIUS, normalized ncDNA (nncDNA) and identification letters for different species. The rows are sorted according to their PIUS values.

| Species | % ncDNA | PIUS | Normalized ncDNA (%ncDNA·PIUS<sup>-1</sup>) | Identifying letter |
| --- | --- | --- | --- | --- |
| *Homo sapiens* | 98.3 | 92000 | 0.0010684783 | A |
| *Mus musculus* | 95 | 62000 | 0.0015322581 | B |
| *Oryza sativa* | 80 | 49000 | 0.0016326531 | C |
| *Arabidopsis thaliana* | 71.2 | 41000 | 0.0017365854 | D |
| *Gallus gallus* | 97 | 30000 | 0.0032333333 | E |
| *Caenorhabditis elegans* | 74.19 | 28000 | 0.0026496429 | F |
| *Drosophila melanogaster* | 81 | 22000 | 0.0036818182 | G |
| *Ciona intestinalis* | 86.8 | 17309 | 0.0050147322 | H |
| *Anopheles gambiae* | 98.3 | 13524 | 0.0072685596 | I |
| *Dictyostelium discoideum* | 43.7 | 12765 | 0.0034234234 | J |
| *Neurospora crassa* | 62.4 | 10200 | 0.0061176471 | K |
| *Streptomyces coelicolor* | 11.1 | 8039 | 0.0013807688 | L |
| *Saccharomyces cerevisiae* | 29.5 | 6050 | 0.0048760331 | M |
| *Pseudomonas aeruginosa* | 10.6 | 5565 | 0.0019047619 | N |
| *Plasmodium falciparum* | 9 | 5449 | 0.0016516792 | O |
| *Schizosaccharomyces pombe* | 42.5 | 5150 | 0.0082524272 | P |
| *Escherichia coli* | 12 | 4400 | 0.0027272727 | Q |
| *Bacillus subtilis* | 80 | 4267 | 0.0187485353 | R |
| *Mycobacterium tuberculosis* | 9 | 3997 | 0.0022516888 | S |
| *Deinococcus radiodurans* | 9.1 | 3085 | 0.0029497569 | T |
| *Fusobacterium nucleatum* | 10.2 | 2046 | 0.0049853372 | U |
| *Neisseria meningitidis* | 17.1 | 2001 | 0.0085457271 | V |
| *Methanocaldococcus jannaschii* | 14 | 1787 | 0.0078343593 | W |
| *Aquifex aeolicus* | 7 | 1553 | 0.004507405 | X |
| *Helicobacter pylori* 26695 | 74.2 | 1553 | 0.0477784932 | Y |
| *Mycoplasma genitalium* | 12 | 483 | 0.0248447205 | Z |

When plotting the data in Table 1 a characteristic graph is obtained (Figure 1). Notably, a direct correlation is not found, but rather organisms with higher PIUS (green) seem to have lower ncDNA percentages than expected. Also, organisms in the other phase of the graph (purple) belong to more primitive phylums. This can be explained by a reduction in nncDNA values in more advanced species.

**Figure 1.**
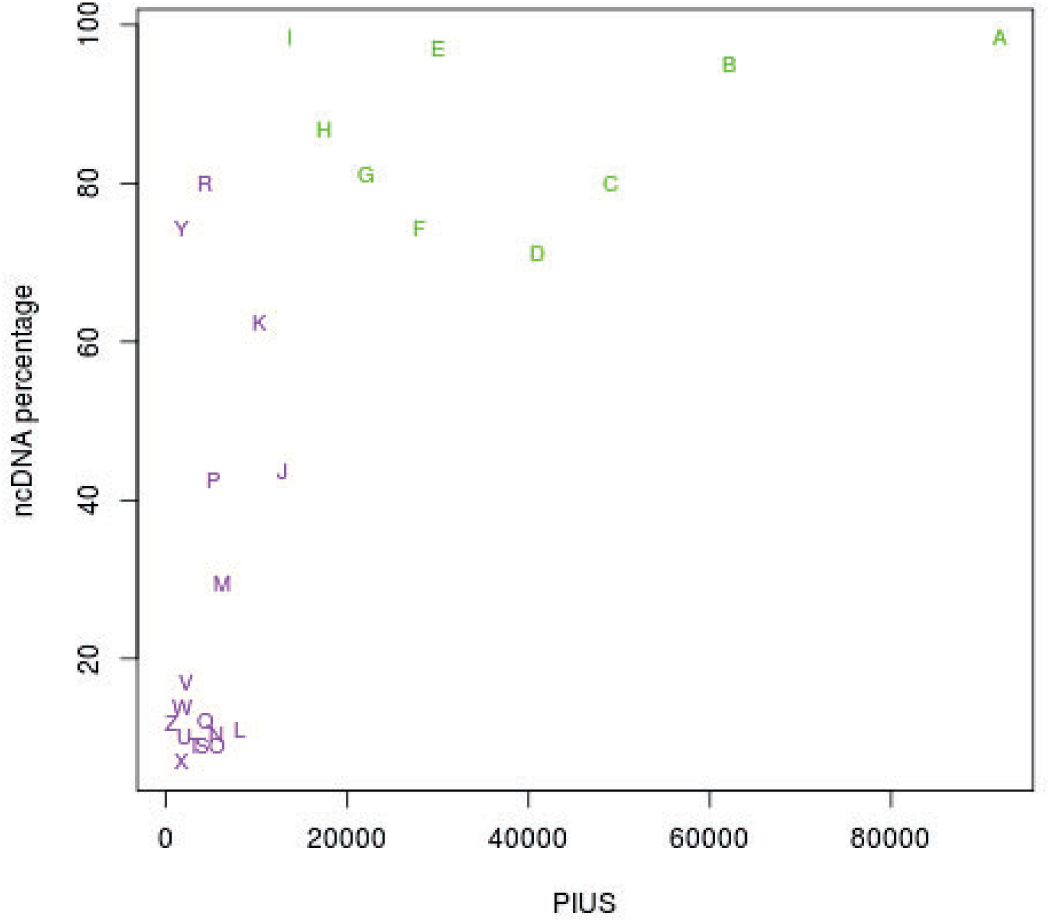
Relation between ncDNA percentage and PIUS for different species. Each organism is represented by a letter as shown in Table 1.

### Normalized ncDNA and control of available complexity

Examination of Figure 1 also reveals a possible limitation on PIUS complexity based on the relation between ncDNA percentage and PIUS. As PIUS values increase, normally so do ncDNA percentages. Since ncDNA percentages cannot logically exceed 100%, there is a limitation on the theoretical maximum PIUS.

Nevertheless, the ratio between ncDNA percentage and PIUS –i.e nncDNA-- is shown to have presumably decreased in evolution. In this manner, more advanced organisms would have achieved greater PIUS by reducing their nncDNA values.

### Exploring the role of chromatin interactions

One of the factors relating ncDNA percentages and PIUS may be regulation of gene expression. At the same time, chromatin interactions are known to regulate gene expression [10]. Therefore, these three elements (ncDNA, PIUS and chromatin interactions) could be connected. To study this possibility, information from the 4DGenome database [11] was retrieved. Particularly, *Mus musculus* and *Homo sapiens* data was selected, as these species are closely related and share most gene regulation mechanisms [12].

The results show that, even though total ncDNA, PIUS and number of chromatin interactions vary substantially from mice to humans, their proportion (k) is approximately constant. This is shown in Table 2:

**Table 2.**
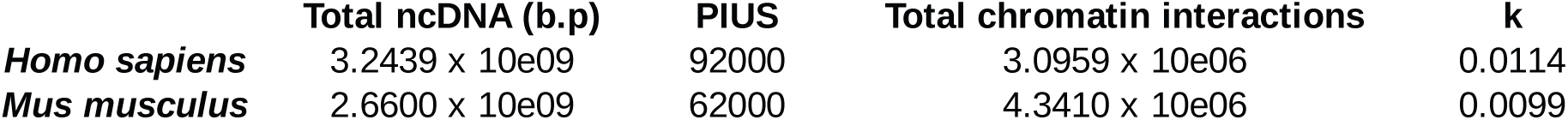
Relation between total ncDNA, PIUS and total chromatin interactions. The parameter k corresponds to the ratio Total ncDNA (b.p) ∙ (PIUS ∙ Total chromatin interactions)⁻¹.

## Discussion

In this work, it is shown that normalized ncDNA has generally decreased in evolution. This is in contrast with ncDNA percentages, where organismal complexity and ncDNA percentages are positively correlated. The advantage of normalized ncDNA over ncDNA percentages is that it includes another complexity indicator, in this case Proteome Information Units.

nncDNA may not be used to measure biological complexity –ncDNA percentages or PIUS are better indicators-- but rather to gauge efficiency in regulation. Since ncDNA contains regulatory DNA, lower nncDNA values indicate more efficient control per Proteome Information Unit.

In addition, lower nncDNAs allow higher theoretical maximum PIUS values. This is in agreement with the observed trend of nncDNA decrease in evolution. However, data (genomic and proteomic) from more species is necessary to confirm these results.

Finally, an empirical constant between ncDNA, PIUS and chromatin interactions is presented for humans and mice. In this manner, PIUS and chromatin interactions seem to be inversely proportional. Information from other related species is also needed to further prove this association.

